# Discovery of integrons in Archaea: platforms for cross-domain gene transfer

**DOI:** 10.1101/2022.02.06.479319

**Authors:** Timothy M. Ghaly, Sasha G. Tetu, Anahit Penesyan, Qin Qi, Vaheesan Rajabal, Michael R. Gillings

## Abstract

Horizontal gene transfer between different domains of life is increasingly being recognised as an important driver of evolution, with the potential to provide the recipient with new gene functionality and assist niche adaptation^1-3^. However, the molecular mechanisms underlying the integration of exogenous genes from foreign domains are mostly unknown. Integrons are a family of genetic elements that facilitate this process within Bacteria via site-specific DNA recombination^4-7^. Integrons, however, have not been reported outside Bacteria, and thus their potential role in cross-domain gene transfer has not been investigated. Here we show that integrons are also present among diverse phyla within the domain Archaea. Further, we provide experimental evidence that integron-mediated recombination can facilitate the recruitment of archaeal genes by bacteria. Our findings establish a new mechanism that can facilitate horizontal gene transfer between the two domains of prokaryotes, which has important implications for prokaryotic evolution in both clinical and environmental contexts.

## Main

Horizontal gene transfer between different domains of life can be a major driver in species evolution^8^. There are now numerous examples of genes that have been transferred between Archaea, Bacteria and Eukarya^3,9-13^. Among the consequences of such gene transfers are the gain of novel biochemical functions and the ability to colonise specific environmental niches^1-3^. However, the molecular mechanisms for most of these transfer events are unknown.

Integrons are genetic elements known to facilitate horizontal gene transfer within Bacteria^4-7^. Integrons can capture exogenous genes, known as gene cassettes, by site-specific recombination. Gene cassette capture is mediated by an integron integrase (IntI), which catalyses the recombination between the recombination site of the inserting cassette (*attC*) and the endogenous integron attachment site (*attI*), immediately adjacent to the *intI* gene. Multiple gene cassettes can be inserted within a single integron, forming cassette arrays that range from 1 to 200+ sequential cassettes^4,6^. Integrons are mostly known for their role in driving the global antibiotic resistance crisis by disseminating diverse resistance determinants among bacterial pathogens^14,15^. However, it is now clear that integrons play a much broader role in bacterial evolution and niche adaptation^16^. The functions encoded by integron gene cassettes are extraordinarily diverse and extend far beyond those of clinical relevance^7,17,18^.

To date, integrons have only been found within bacterial genomes, where they have been detected within diverse phyla^19^. However, gene cassette amplicon sequencing has yielded cassette-encoded proteins that share homology with archaeal proteins^20,21^. Without broader genomic context, however, the taxonomic residence of such gene cassettes is unknown.

Here, we screened all publicly available archaeal genomes to show for the first time that integrons are not limited to Bacteria, but are also present in Archaea. Archaeal integrons exhibit the same characteristics and functional components as bacterial integrons. Further, we demonstrate experimentally that diverse archaeal gene cassettes can be successfully recruited by a bacterial host, facilitated by integron-mediated recombination. Such a mechanism can potentially facilitate a cross-domain highway of gene transfer between Archaea and Bacteria, with important implications for prokaryotic evolution.

### Discovery of integrons in Archaea

Here, we report the discovery of integrons in the domain Archaea. We screened 6,718 archaeal genomes for integrons using the standard criteria applied to integron surveys in Bacteria^19,22,23^. These include the presence of integron integrase genes and/or clusters of gene cassette *attC*s (defined as at least two *attC*s with less than 4 kb between each). We identified integrons in 75 archaeal metagenome-assembled genomes (MAGs) from 9 phyla (Fig. 1 and Supplementary Table 1). It is not surprising that integrons were detected only in MAGs, given that they constituted ∼95% of all available archaeal genomes. However, to ensure that these integrons did not arise from contaminating bacterial contigs, incorrectly binned with archaeal MAGs, we applied stringent MAG refinement and quality filtering (see Methods for details). Additionally, we found that ∼7% of integron-bearing MAGs had at least one archaeal phylogenetic marker gene on the same contig as an integron (Supplementary Table 2), confirming these to be located on archaeal chromosomes. No integron was ever co-located with a bacterial marker gene. The markers used for this analysis consisted of a comprehensive set of 122 archaeal and 120 bacterial proteins identified as suitable for phylogenetic inference^24^.

**Fig. 1:**
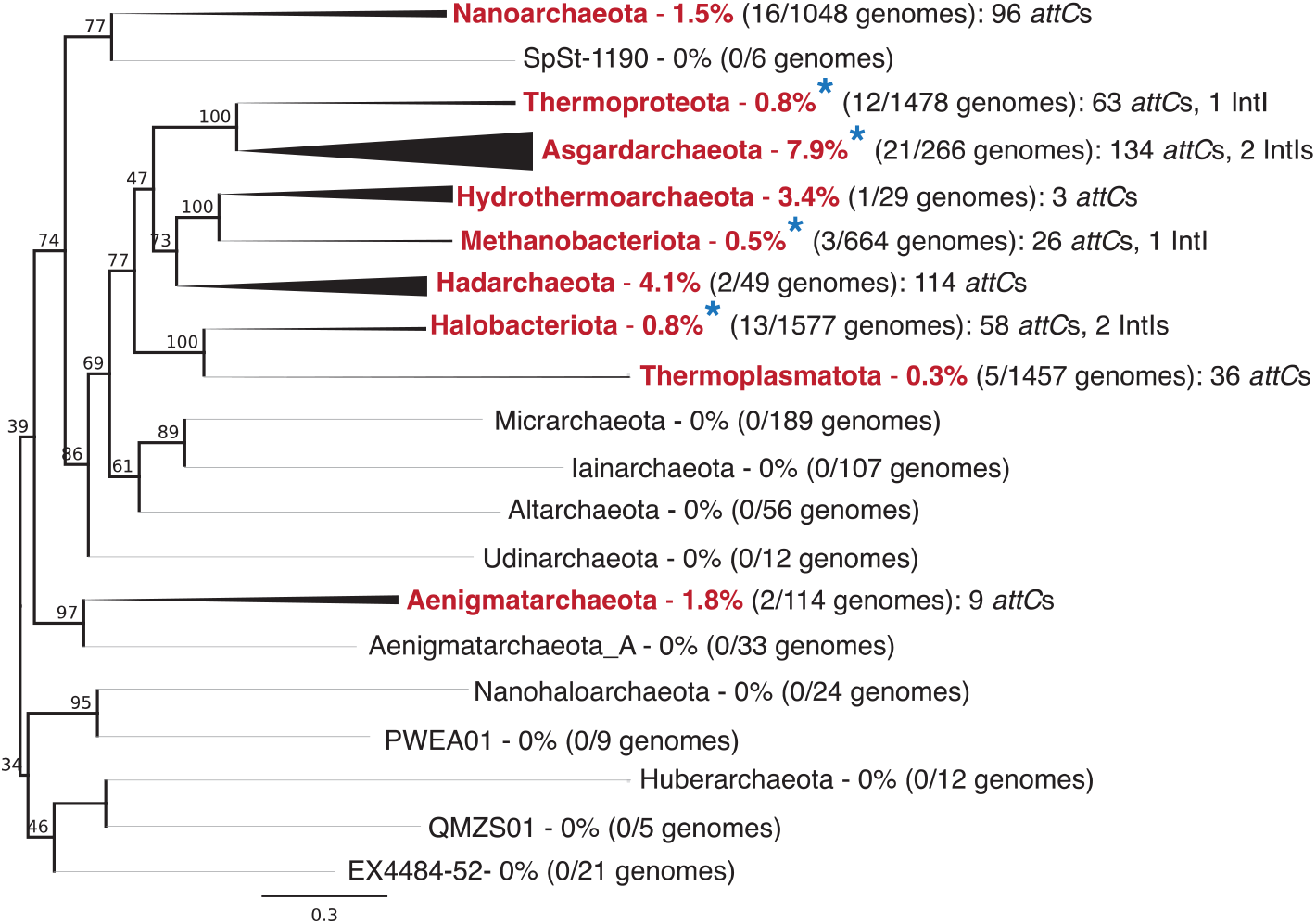
Phylogenetic distribution of integrons among Archaea. Archaeal phyla found to carry integrons are labelled in red, and those found to have an integron integrase gene (*intI*) are denoted with blue asterisks. Branch thickness indicates the proportion of genomes with integrons for each phylum.

Among the 75 archaeal genomes, we detected six IntIs and 539 *attC* sites (excluding all singleton *attC*s). We found that archaeal *attC*s and IntIs are largely restricted to one clade of Archaea (Fig. 1), with some outliers, suggesting that integron diversification, for the most part, has likely occurred within one archaeal clade, with occasional horizontal movements to other archaeal phyla. In particular, integrons were significantly enriched in the phylum Asgardarchaeota (χ^2^ test, p < 0.00001) (Fig. 1), being detected in almost 8% of available Asgard genomes. Asgardarchaeota contributed the most genomes with detectable integrons (28%) and the greatest number of gene cassettes (24.9%), despite having relatively few genomes among the dataset (comprising 4% of available archaeal genomes). We also detected integrons in 3-4% of genomes from the phyla Hadarchaeota and Hydrothermoarchaeota (Fig. 1), although these comprised few available genomes (n<50). A skewed phylogenetic distribution of integrons has similarly been observed among Bacteria^19^. For example, in the phylum Proteobacteria, integrons are enriched within the class Gammaproteobacteria (20% of genomes), while being entirely absent from its sister class Alphaproteobacteria. This is intriguing given that integrons have been detected at widely varying prevalence in more distantly related bacterial phyla such as Cyanobacteria, Spirochaetota, Planctomycetota, Chloroflexota, Bacteroidota and Desulfobacterota^19,22^.

### Genetic structure of archaeal integrons

We found that archaeal integrons exhibit the same structure and functional components as bacterial integron cassette arrays (Extended Data Fig. 1). That is, tandem arrays of short open reading frames (ORFs), generally in the same orientation, interspersed by *attC* recombination sites. Archaeal *attC*s exhibit the same single-stranded folding structure as bacterial *attC*s, which is essential for them to act as structure-specific DNA recombination sites^25-31^. We also note that archaeal IntIs exhibit the defining characteristics of bacterial IntIs, being tyrosine recombinases that possess a unique IntI-specific additional domain surrounding the patch III motif region necessary for integron-mediated recombination^32^. We found examples of ‘complete’ integrons, these being cassette arrays adjacent to a detectable *intI* gene (Extended Data Fig. 1). We also found examples of putative *attI* sites, which act as insertion points for incoming gene cassettes. These *attI*s were immediately downstream of the *intI* gene, semi-conserved across distinct archaeal phyla (Extended Data Fig. 2a,b), and exhibited the same canonical insertion point as all known bacterial *attI*s (Extended Data Fig. 2c).

Most archaeal integrons that we identified were CALINs (clusters of *attC*s lacking integron integrases; Supplementary Table 3). This is not surprising given the fragmented nature of MAGs, and the high prevalence of CALINs also found in bacterial genomes. Indeed, among Bacteria, CALINs are more abundant than complete integrons that possess an *intI* gene, and exhibit a much wider taxonomic distribution^19^. Two In0 elements were also detected among Archaea. These are integrons that have an *intI* gene without an adjacent *attC* site (Extended Data Fig. 1). However, both archaeal genomes with an In0 also had clusters of *attC* sites on other contigs. Among our dataset, the longest array of *attC*s on the same contig was 12, however, we found as many as 107 *attC*s (over 18 contigs) within a single MAG (Supplementary Table 1). The number of *attC*s within a single MAG ranged from 2 to 107, with an average of 7 *attC*s.

### Platforms for cross-domain gene transfer

Archaeal gene cassettes with *attC*s from diverse phyla can be recognised and recruited by Bacteria (Fig. 2). We demonstrate that cassette insertion (*attC* x *attI* recombination) can occur following the conjugation of circular DNA molecules with archaeal *attC*s into an *Escherichia coli* recipient harbouring a bacterial class 1 integron (Fig. 2a). Insertion events were confirmed with Sanger sequencing of the PCR-amplified *attC*/*attI* recombination junctions (Fig. 2a, Extended Data Fig. 3). We found that recruitment of cassettes with archaeal *attC*s occurred at similar frequencies to that of the paradigmatic bacterial *attC* site, *attC*_*aadA7*_, which we used as a positive control (Fig. 2b, Extended Data Table 1). We observed an average recombination frequency of 2.5×10^−1^ between *attI1* and *attC*_*aadA7*_. Comparable frequencies (ranging from 1.9×10^−4^ – 3.2×10^−1^, with an average of 5.1×10^−2^) were observed for eight out of nine archaeal *attC*s (Kruskal-Wallis test, p=0.488), which were selected from multiple archaeal phyla. Further, we confirmed that cassette recruitment was mediated by IntI1 activity, since no *attC* x *attI* recombination events were detected when *intI1* was absent or when its expression was suppressed (Extended Data Table 1). We therefore show that integron-mediated gene transfer can occur between the two domains of prokaryotes.

**Fig. 2:**
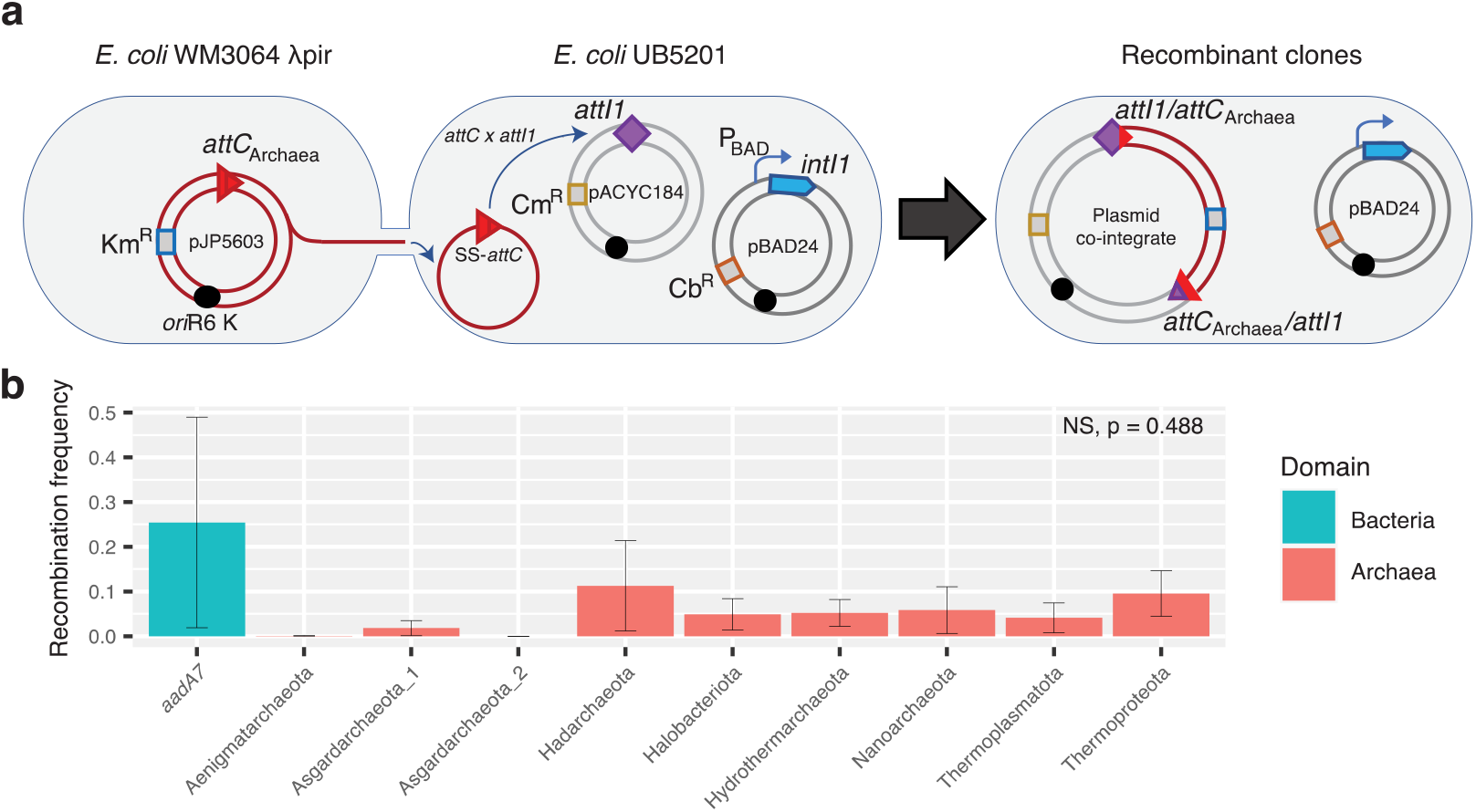
Cassette recruitment (*attC* x *attI* recombination) assays. **a**, schematic outlining the experimental setup of the cassette insertion assays. The suicide vector pJP5603 with an *attC* site is delivered into the recipient *E. coli* UB5201 strain via conjugation. The recipient strain carries an *intI1* gene, expressed from the inducible P_BAD_ promoter, and an *attI1* site, residing on the pBAD24 and pACYC184 backbones, respectively. The donor suicide vector cannot replicate within the recipient host, and thus, can only persist following *attC* x *attI* recombination to form a plasmid co-integrate. **b**, average recombination frequencies (±1 S.E.) between *attI1* and nine archaeal *attC*s (with phyla of origin labelled along the X-axis) and the paradigmatic bacterial *attC* site (*attC*_*aadA7*_), used as positive control. Average frequencies were calculated following three independent cassette insertion assays (see Methods for details). No statistically significant difference in recombination frequencies were detected among the tested *attC*s (Kruskal-Wallis test, n=27, df=8, p=0.488). Recombination frequencies are shown for *attC* bottom strands only. See Extended Data Table 1 for *attC* top strand recombination frequencies.

Importantly, we find that the most clinically significant class of integrons (class 1) can recruit archaeal cassettes as efficiently as bacterial cassettes. Class 1 integrons are highly promiscuous due to their association with diverse mobile genetic elements, facilitating their spread into at least 100 bacterial species^7^. They collectively carry more than 130 different resistance genes^14^, most of which are of unknown taxonomic origin^22^. Our findings open the possibility that Archaea could be an unexplored source of class 1 integron gene cassettes.Regardless, our findings indicate that any bacterial strain with a class 1 integron has the capacity to incorporate exogenous genes from diverse archaeal phyla, greatly expanding the genetic pool that they have access to.

The cross-domain transfer of integron gene cassettes is possibly widespread. For example, we detected 23 *attC*s from six archaeal genomes that exhibited 95-100% nucleotide identity to *attC*s within sequenced bacterial integrons (Supplementary Table 4). The archaeal *attC*s were from three phyla: Nanoarchaeota, Thermoproteota and Hadarchaeota. The homologous *attC*s in Bacteria were found in 26 genomes from 5 phyla: Proteobacteria, Spirochaetota, Myxococcota, Nitrospirota and Desulfobacterota. One of these *attC* sites was associated with a class 1 integron gene cassette, encoding an NADPH-dependent oxidoreductase found on five different Enterobacteriaceae plasmids (Supplementary Table 4). In Archaea, however, this *attC* site was part of a cassette that encoded a ligand-binding protein of unknown function. Nevertheless, since strong *attC* homology is a characteristic of cassettes that share the same taxonomic origin^22,33,34^, it is possible that some clinically relevant gene cassettes now found on class 1 integrons might be of archaeal origin.

### Diversity of archaeal integrons

#### Diversity of integron integrases

Archaeal IntIs are phylogenetically distinct from bacterial IntIs (Fig. 3). We detected six IntIs from four archaeal phyla (Fig. 1), however, three of these were excluded from further phylogenetic analysis based on either short sequence length (< 200 amino acids) or partial coverage of the IntI-specific domain (Extended Data Fig. 4). We found that archaeal IntIs form their own monophyletic clade separate from known bacterial IntIs^22^. This strongly suggests that IntI radiation has occurred within Archaea and that their distribution, at least among the archaeal genomes in our dataset, is not likely to be the result of multiple IntI acquisitions from Bacteria. Regardless, we show that IntIs from distinct archaeal phyla, isolated from different environments, are more closely related to each other than they are to any bacterial IntI.

**Fig. 3:**
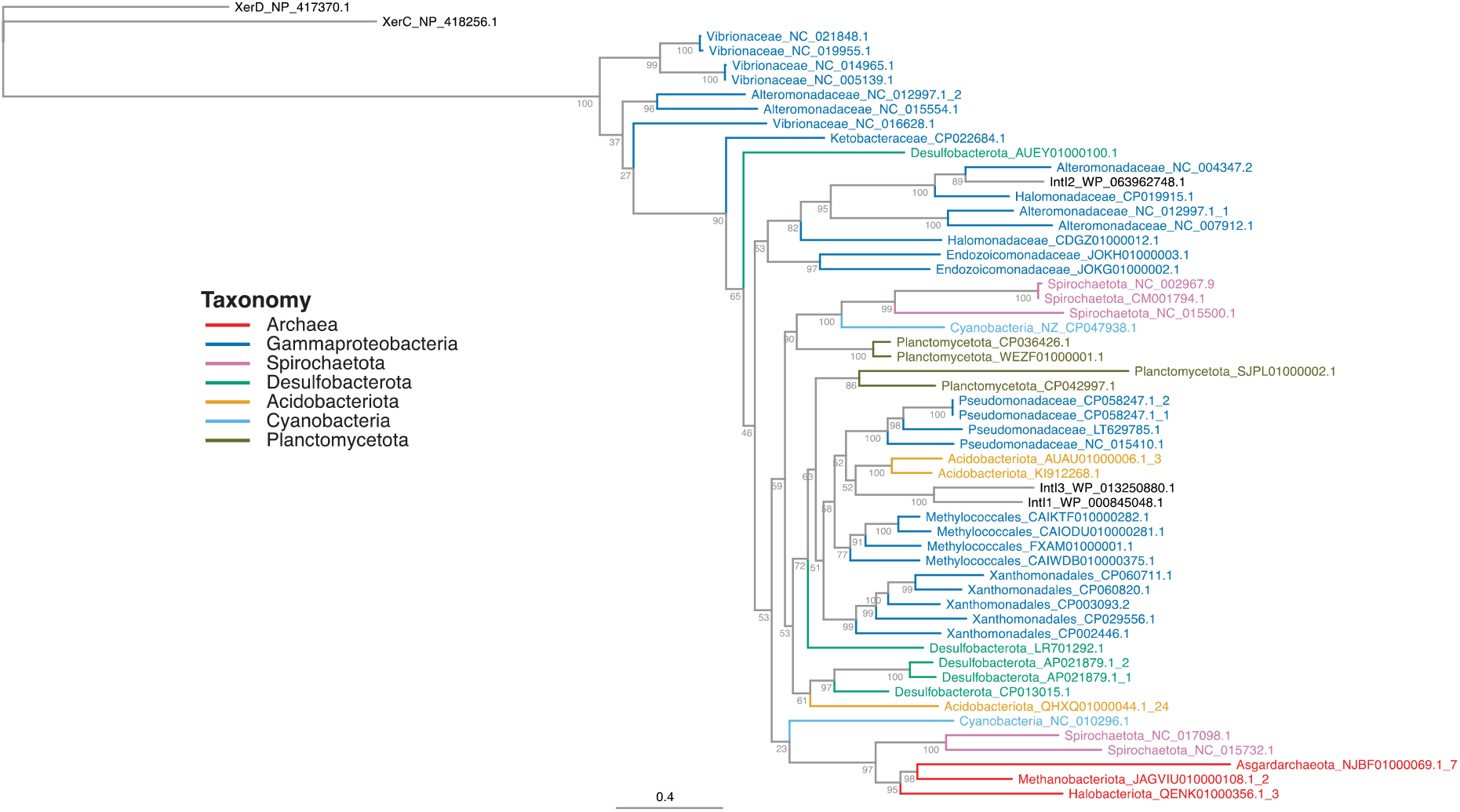
Phylogeny of integron integrases from Archaea and Bacteria. To root the tree, the tyrosine recombinases XerC and XerD from *Escherichia coli* were used as outgroups. Integron integrases (IntIs) are coloured according to their taxonomy.

The closest sister clade to the archaeal IntIs comprises two Spirochaetota IntIs (Fig. 3). Intriguingly, these two IntIs are phylogenetically distinct from ‘typical’ Spirochaetota IntIs, which are generally in reverse orientation^5,35^. Further, the two Spirochaetota that harboured atypical IntIs were isolated from extreme environments: a brine layer within an alkaline lake and a hot spring, respectively; environments known to have a relatively high abundance of Archaea^36^. Thus, these atypical Spirochaetota IntIs might have been horizontally acquired from Archaea that share the same extreme environments.

#### Diversity of attC recombination sites

Archaeal *attC*s exhibit broad sequence and structural diversity (Fig. 4a). We find that some archaeal phyla possess *attC*s with a restricted diversity (e.g., Hadarchaeota and Aenigmatarchaeota), while other phyla have extremely variable *attC*s distributed throughout the *attC* diversity space (e.g., Asgardarchaeota, Nanoarchaeota and Thermoproteota). This distribution could indicate that different taxa have different propensities for horizontal exchange of gene cassettes^7,22^. We show that archaeal *attC*s are significantly more similar within a genome than between genomes (Fig. 4b). This characteristic is also a hallmark of chromosomal bacterial integrons^19,34^. We also show that *attC*s are more similar between different genomes from the same archaeal order than they are between genomes from different orders (Fig. 4c). This order-level *attC* homology is also seen within Bacteria^22,33^. Thus, the ecological and evolutionary forces that promote and/or constrain *attC* diversity^7^ are likely to be similar for both Archaea and Bacteria.

**Fig. 4:**
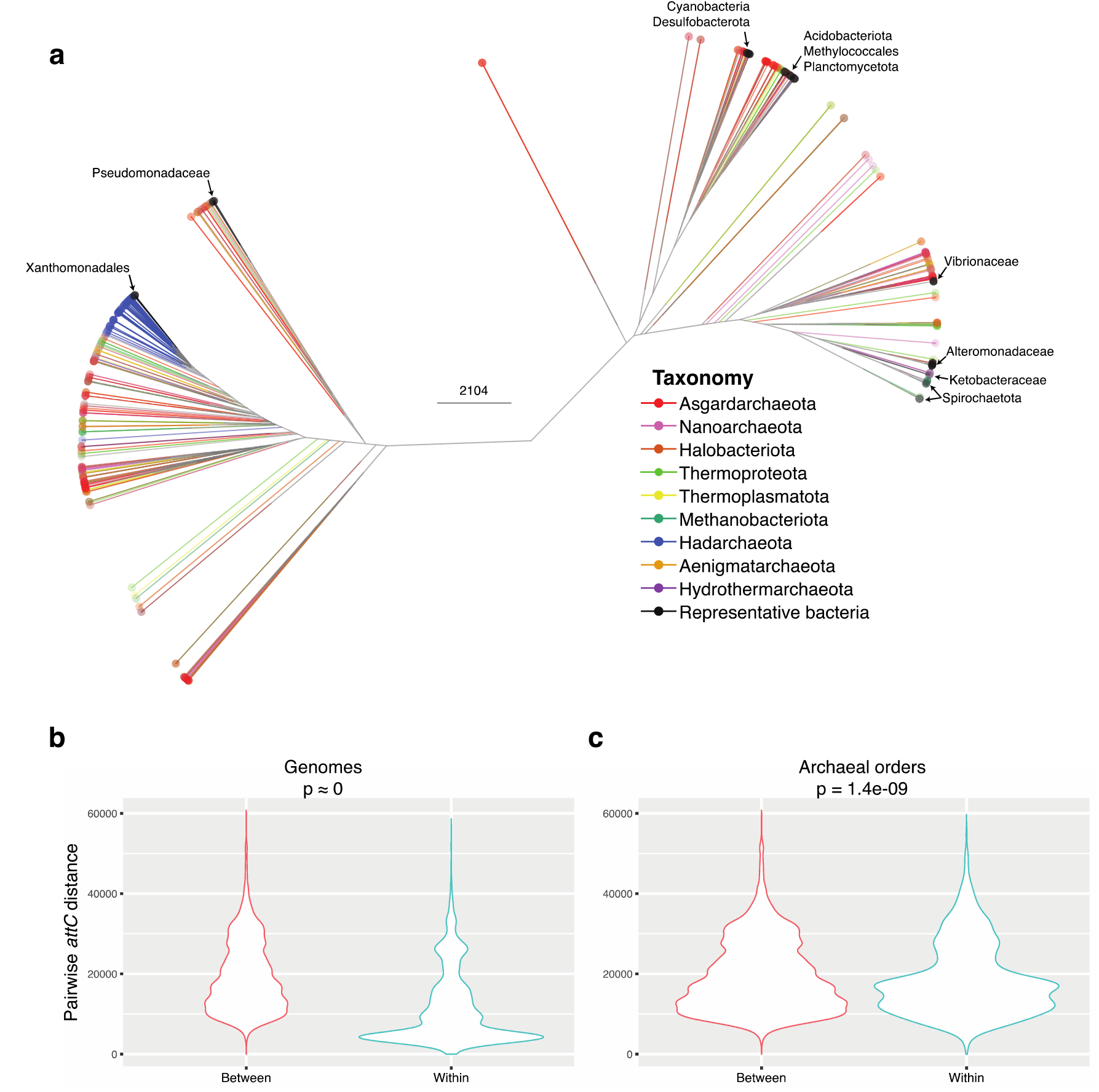
Structural and sequence diversity of archaeal *attC* recombination sites. **a**, structure-based clustering of all archaeal and representative bacterial *attC*s. Branches and tips are coloured according to archaeal phylum. The taxa of bacterial *attC*s are labelled with arrows. **b**, distribution of the sequence and structural distances calculated for all pairwise comparisons of *attC*s within and between genomes. **c**, distribution of distances for all pairwise comparisons of *attC*s from different genomes that are either from the same or different archaeal orders.

There is a clear overlap in the sequence and structural diversity of *attC*s from Archaea and Bacteria (Fig. 4a). This provides additional evidence that the mechanistic overlap between archaeal and bacterial *attC*s is extensive, and thus, cross-domain transfer of cassettes could be common in shared environments. It also suggests that the recruitment of extra-domain gene cassettes can be facilitated by diverse classes of integrons, of which there are thousands (based on IntI amino acid homology^37^). The broad distribution of integrons among the two domains suggest that integron-mediated transfer plays an important role in prokaryotic evolution.

#### Functional diversity of gene cassettes

We detected 549 cassette-encoded proteins among Archaea. Only 23.1% of these could be classified into a known COG category (Extended Data Fig. 5). In contrast, 47.4% of all proteins from the 75 integron-bearing archaeal genomes could be assigned a known COG category. This underrepresentation (χ^2^ test, p < 0.00001) of known COGs among cassette proteins has previously been reported for bacterial integrons^4,5,33^. To gain further insight into possible cassette functions, eggNOG 5.0^38^ and Pfam^39^ database searches were performed, assigning putative functions to 228 (41.5%) of the archaeal cassette-encoded proteins. Out of those with functional predictions, proteins involved in toxin-antitoxin (TA) systems (10.5%); phage resistance proteins via DNA methylation or restriction endonuclease activities (8.3%); and acetyltransferases (4.4%) were particularly prevalent (Supplementary Table 5). These are the functions most commonly reported for gene cassettes in Bacteria^5,7,33,34,40^. TA gene cassettes are particularly common in bacterial integrons, where they can stabilise very large cassette arrays^41,42^. The antitoxin modules of TA cassettes can also counteract the toxins of homologous systems found on plasmids and phage, thus potentially protecting their host from invading mobile elements^43,44^.

In addition, 13.2% of archaeal cassette-encoded proteins had signal peptides, which represents a significant enrichment relative to their broader genomic contexts (6.9%, χ^2^ test, p < 0.00001). Signal peptides are short amino acid tag sequences that target proteins into, or across, membranes. Again, transmembrane and secreted proteins are commonly encoded by gene cassettes in Bacteria^33^, and are hypothesised to help facilitate interactions with their broader environment^7^.

Indeed, we find that functions of archaeal cassettes are associated with their environment (Fig. 5). Functional families cluster according to their specific environment, and these environmental clusters, in turn, group according to their broader environmental type (Fig. 5). This environmentally explicit clustering might be the result of local ecological and evolutionary forces. That is, gene cassettes in Archaea confer niche-specific functional traits and/or horizontal transfer of cassettes occurs between archaeal phyla co-located in the same environment.

**Fig. 5:**
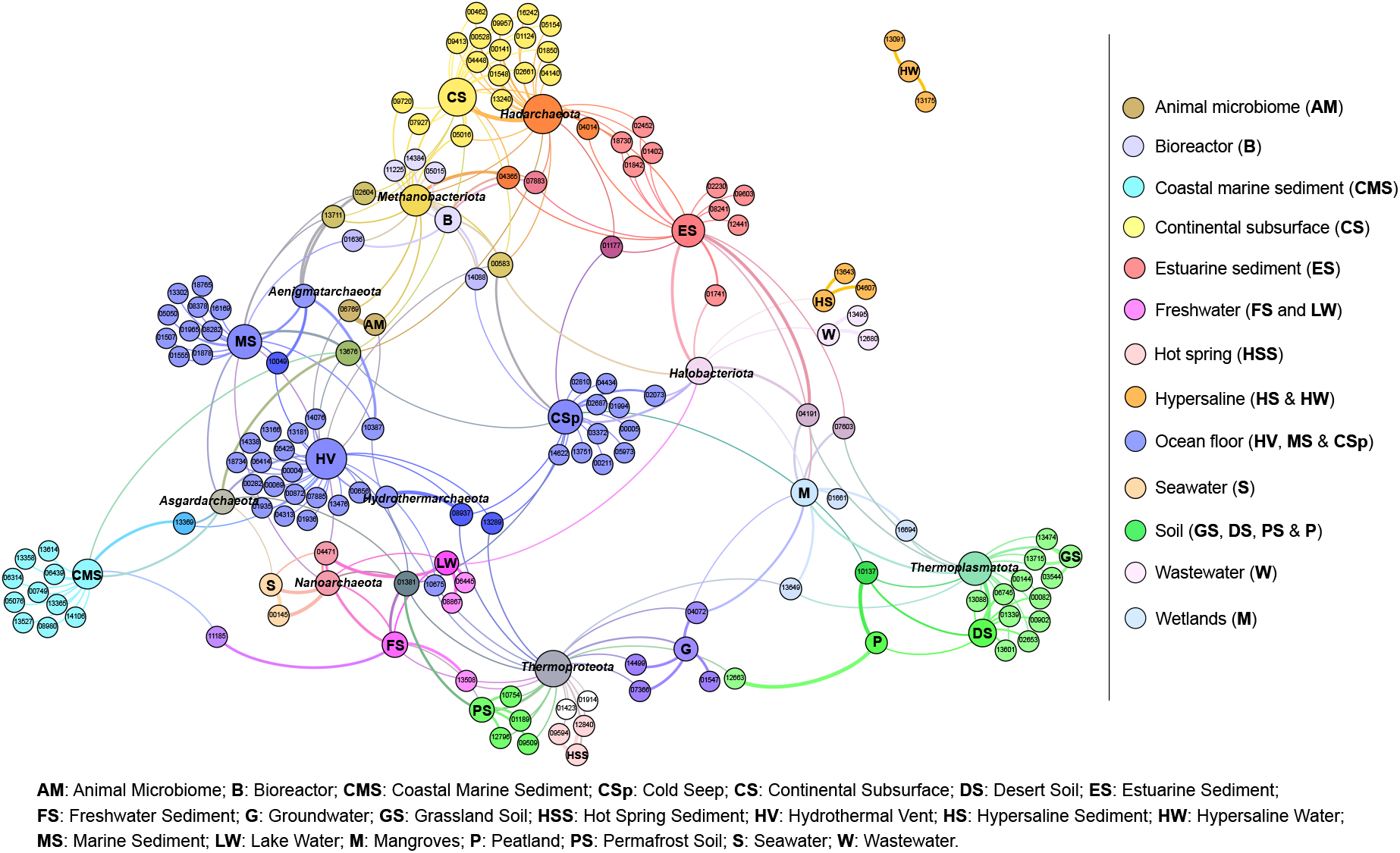
A network linking Pfam functions of archaeal integron gene cassettes with their taxonomic and environmental contexts. The force-directed representation of the network is constructed based on co-occurrence patterns and correlations (p < 0.05) between Pfam functions, taxonomic groups, and specific environments from which the organisms were sampled. Nodes that represent taxonomic groups and specific environments are labelled accordingly. All other nodes denote Pfam functions and are labelled with a Pfam number preceded by ‘PF’. Specific environments are grouped into broader environment types, each of which is coloured as per the panel. Pfams directly linked to specific environment types are coloured in corresponding colours. Pfams linked to more than one environment type are coloured in overlapping colours. The size of the node is relative to the node authority based on the degree of correlations. Edges (the lines connecting the nodes) represent correlations between nodes. Edge colour denotes the overlapping colour of the two nodes it connects. Edge thickness represents the strength of correlation. The full description of all correlations and Pfam functions is presented in Supplementary Table 6.

## Conclusion

Here, we present the first evidence of integrons in the domain Archaea. We demonstrate that they have the same functional characteristics as bacterial integrons. We also present experimental evidence that bacteria can successfully recruit archaeal gene cassettes, facilitated by integron-mediated DNA recombination. Our results thus establish a novel mechanism for cross-domain gene transfer between Archaea and Bacteria. We also find that, although archaeal IntIs are phylogenetically distinct from bacterial IntIs, their associated *attC* recombination sites are shared with Bacteria. This suggests that integron-mediated cross-domain gene transfer is widespread and plays an important role in prokaryotic evolution.

## Methods

### Data acquisition and quality filtering

All available archaeal genomes were downloaded from the NCBI Assembly Database (n=8,160; last accessed 2021-Oct-5). Of these, ∼ 95% were metagenome-assembled genomes (MAGs). We applied stringent filtering criteria to remove low quality MAGs. First, to improve MAG quality, we identified and removed contaminating contigs from each MAG using MAGpurify v2.1.2^45^ with the following modules: ‘*phylo-markers’*, which finds taxonomically discordant contigs using 100 archaeal and 88 bacterial single-copy taxonomic marker genes from the PhyEco database; ‘*clade-markers’*, which finds contaminating contigs using a database of 855,764 clade-specific prokaryotic marker genes (MetaPhlAn2 database^47^); ‘*tetra-freq’*, which employs principal component analysis (PCA) to identify contaminating contigs with outlier tetra-nucleotide frequency; and ‘*gc-content’*, which uses PCA to identify contaminating contigs with outlier GC content.

After refinement, the quality of the genomes was assessed using CheckM v1.1.3^48^, which uses single-copy lineage-specific marker genes to estimate genome completeness and contamination. There is strong community consensus that high quality MAGs are those that are more than 90% complete and have less than 5% contamination, while medium quality MAGs have a completeness ≥50% and contamination <10%^24,45,49-52^. In this context, however, we were more concerned with the level of contamination than completeness, and thus removed all genomes with an estimated contamination ≥ 5%. The completeness of the remaining genomes ranged from 15% – 100%, with a median of 81%. The estimated contamination ranged from 0% – 4.98%, with a median of 0.93%.

Archaeal genomes were assigned taxonomic classifications based on the Genome Taxonomy Database (GTDB)^49-51^ using GTDB-Tk v1.6.0^53^ with release 06-RS202 of the GTDB. We employed the *classify_wf* command with default settings. This workflow identifies and aligns 120 bacterial and 122 archaeal phylogenetic marker genes^24^. GTDB-Tk then classifies each genome based on its placement into domain-specific reference trees (built from 47,899 prokaryote genomes), its relative evolutionary divergence, and average nucleotide identity to reference genomes in the GTDB. Any genomes not classified within the domain Archaea were removed. This resulted in a final set of 6,718 archaeal genomes retained for further analysis.

To infer the phylum-level phylogeny of Archaea, the highest quality representative genome from each phylum was selected based on its genome quality score (defined by Parks et al.^24^ as the estimated completeness of a genome minus five times its estimated contamination). From representative genomes, a concatenated multiple protein sequence alignment of the 122 archaeal phylogenetic markers was generated using GTDB-Tk v1.6.0^53^. A maximum-likelihood tree was generated from the alignment using IQ-TREE v1.6.12^54^ with the best-suited protein model as determined by ModelFinder^55^ and 1,000 bootstrap replicates [parameters: -m MFP -bb 1000].

### Integron detection

Due to faster processing speeds of large datasets, we initially screened all filtered genomes for *attC* recombination sites using *attC*-screening.sh^37^(https://github.com/timghaly/integron-filtering) with default parameters. This script uses the HattCI^56^ + Infernal^57^ pipeline (first described by Pereira *et al*.^23^) to search for the conserved sequence and structure of *attC* sites. Genomes that had at least one detectable *attC* site were additionally screened using IntegronFinder v2.0rc6^19^ [parameters: --local-max --cpu 24 -- gbk], which searches for integron integrases and gene cassette arrays. Only IntIs, *attCs* and cassette-encoded proteins identified by IntegronFinder were included in downstream analyses.

To ensure that these integrons were not from contaminating bacterial contigs that had been incorrectly binned with archaeal MAGs, we screened all contigs containing an integron for prokaryotic marker genes using GTDB-Tk v1.6.0^53^. These consisted of 122 archaeal and 120 bacterial proteins identified as suitable phylogenetic markers^24^. We found a total of nine prokaryotic marker genes among seven integron-bearing contigs. All nine markers were confirmed to be archaeal via a BLASTP search of the NCBI nr database (Supplementary Table 2).

### Analysis of integron integrases, attC sites and cassette-encoded proteins

IntegronFinder identifies IntIs using the overlap of two protein hidden Markov model (HMM) profiles. The first is the Pfam profile PF00589 to identify tyrosine recombinases, and the second is a protein profile built from the IntI-specific domain that separates IntIs from other tyrosine recombinases^32^. Identified archaeal IntIs, with matches to both protein profiles, were placed in a phylogeny alongside a set of previously identified bacterial IntIs^22^. IntIs shorter than 200 amino acids or those that did not span the complete IntI-specific domain were removed from phylogenetic analysis. The remaining IntIs were aligned using MAFFT v7.271^58^ [parameters: --localpair --maxiterate 1000] and trimmed using trimAl v1.2rev59 [parameters: -automated1]. A maximum-likelihood tree was generated from the alignment using IQ-TREE v1.6.12^54^ with the best-suited protein model as determined by ModelFinder^55^ and 1,000 bootstrap replicates [parameters: -m MFP -bb 1000].

The sequence and structural diversity of *attC*s was assessed using RNAclust v1.3^59^ as previously described^22^. RNAclust uses LocARNA^60,61^ to generate pairwise structural alignments (based on both sequence and folding structure) of input sequences. RNAclust then calculates pairwise distances to create a hierarchical-clustering tree from a WPGMA analysis. All archaeal *attC*s along with a set of previously identified *attC*s from representative bacterial taxa^22^ were clustered using RNAclust’s default parameters.

Cassette-encoded proteins identified by IntegronFinder were functionally annotated using InterProScan v5.44-79.0^62^, with default parameters against the Pfam^39^ database, and eggNOG-mapper v2.0.1b^63,64^, executed in DIAMOND^65^ mode against the eggNOG 5.0 database^38^. To identify cassettes that encode transmembrane and secreted proteins, we searched protein sequences for prokaryotic signal peptides using SignalP 5.0^66^ with default parameters. The correlation analysis of cassette functions was performed as described in Penesyan et al^67^. Briefly, Pearson’s correlations, based on co-occurrences between Pfam functions, specific environments and archaeal phyla were calculated using the Hmisc v4.5-0 R package^68^. The network was generated from all positive correlations with p-values <0.05 using the ForceAtlas2 layout algorithm^69^ within the Gephi software^70^. Specific correlations and the description of Pfam functions are listed in Supplementary Table 6.

### Bacterial strains and plasmids for attC recombination assays

The bacterial strains and plasmids used in this study are listed in Supplementary Table 7. LB medium (Lennox) was used to grow bacterial strains supplemented with appropriate antimicrobial agents. The final concentrations of antimicrobial agents used were kanamycin (Km) = 50 µg/mL, carbenicillin (Cb) = 75 µg/mL, and chloramphenicol (Cm) = 20 µg/mL. LB medium was supplemented with 0.3 mM 2,6-diaminopimelic acid (DAP) to culture the auxotrophic *E. coli* WM3064 λpir strain^71^.

### Construction of attC donor strains

Nine archaeal *attC*s, selected from diverse archaeal phyla (Supplementary Table 8) along with one bacterial *attC* (*attC*_*aadA7*_) were used for the recombination assays. Two donor strains were constructed for each *attC*, delivering either the *attC* top or bottom strands via conjugation. Overlapping forward and reverse primers were designed to generate each *attC* sequence flanked by *Xba*I and *Bam*HI overhangs respectively (e.g. primer pair *attC*-*aadA7*-FW/REV for *attC*_*aadA7*_). The annealed primer dimers were then ligated into the mobilisable suicide vector pJP5603^72,73^. The *attC* top strand donor strains were generated by transforming the ligation product into electrocompetent cells of the DAP auxotrophic *E. coli* strain WM3064 λpir. Using the same procedures, all *attC* top strand donor plasmids and strains were constructed using the pairs of long primers listed in Supplementary Table 9.

To deliver *attC* bottom strands, the pJP5603rev (pJPrev) vector was generated to invert *oriT* orientation relative to that of the pJP5603 parental vector. The multiple-cloning site and vector backbone of pJP5603 were PCR amplified using the primer pairs pJP-MCS-FW/REV and pJP-Backbone-FW/REV respectively (with *Xho*I and *Mlu*I restriction sites introduced) followed by restriction digest and ligation. The same primer pairs for generating the top strand donor plasmids were used to create the bottom strand donor plasmids and strains by cloning the same *attC* sequences into the *Xba*I/*Bam*HI sites of pJPrev.

### Construction of the recipient strain

We generated a recipient strain using *E. coli* UB5201^74^ that carried the *intI1* gene and the *attI1* recombination site residing on the pBAD24^75^ and pACYC184^76^ backbones, respectively. The *intI1* gene of the R388 plasmid^77^ was PCR amplified using the primer pair *intI1_Eco*RI-F/*intI1*_*Hin*dIII-R (Supplementary Table 9). The L-arabinose inducible pBAD::*intI1* plasmid was generated by cloning *intI1* into the pBAD24 expression vector. The pACYC184::*attI1* recipient plasmid was created by assembling the *attI1* sequence (from R388) into the pACYC184 plasmid backbone using the NEBuilder HiFi DNA Assembly Cloning Kit (New England Biolabs, United States). The PCR products required for the assembly were generated using the *attI1*_fw/*attI1*_rev and pACYC184_backbone_F/pACYC184_backbone_R primer pairs. *E. coli* UB5201 strain was co-transformed with pBAD::*intI1* and pACYC184::*attI1* to generate the *E. coli* UB5201 + pBAD::*intI1* + pACYC184::*attI1* recipient strain for *attC* x *attI* suicide conjugation assays. *E. coli* UB5201 + pBAD24 + pACYC184::*attI1* was created as an *intI1*-negative control strain. All plasmid constructs were confirmed by Sanger sequencing and restriction enzyme digests.

### attC x attI suicide conjugation assays

The frequencies of recombination between the archaeal *attC* sequences and the class 1 integron *attI1* site were quantified using previously established *attC* x *attI* suicide conjugation methods^25,29,31,78,79^ with minor modifications. Briefly, the Cb-resistant UB5201 + pBAD::*intI1* + pACYC184::*attI1* recipient strain was filter-mated with Km-resistant WM3064 λpir *attC* donor strains in DAP-supplemented LB media. The expression of *intI1* was either induced using L-arabinose (2 mg/mL) or suppressed with D-glucose (10 mg/mL). After 6 hours of incubation at 37°C, the recovered conjugation mix was plated on DAP-free LB agar with Km, as well as on LB agar containing Cb. This method allowed for negative selection of the donor strain, which cannot grow in the absence of DAP, and positive selection of the recombinant recipient clones, which become Km-resistant following plasmid co-integration (Fig. 2a). The recombination frequency was determined as the ratio of the colony forming units (CFU) for Km-resistant recombinants to the CFU for the total number of Cb-resistant recipients after two days of incubation. All assays were performed in three biological replicates, and recombination frequencies were calculated as the mean of the three independent experiments. To confirm the co-integrates, colony PCR was performed on eight randomly chosen colonies per conjugation set for each biological replicate using the following primer pairs pACYC_F/M13F and pACYC_R/M13R (Extended Data Fig. 3). Sanger sequencing of PCR products was performed for four recombinant colonies per conjugation set.

## Data availability

All genome sequences were downloaded from the NCBI Assembly Database (https://www.ncbi.nlm.nih.gov/assembly; last accessed 2021-Oct-5).

## Code availability

All code and software used in this study are described within the manuscript.

## Acknowledgements

The authors would like to thank Ian Paulsen for comments on earlier versions of the manuscript. This work was funded by the Australian Research Council (Discovery Project DP200101874).

## Author contributions

TMG contributed to the conception of the study, performed all data analyses, wrote the original draft of the paper, and contributed to the final editing of the paper. SGT contributed to the conception of the study and the final editing of the paper. AP performed the correlation analysis of cassette functions, and contributed to the final editing of the paper. QQ was involved with the design and implementation of the experimental work, and contributed to the final editing of the paper. VR was involved with the design and implementation of the experimental work, and contributed to the final editing of the paper. MRG contributed to the conception of the study and the final editing and revision of the paper. All authors contributed to the article and approved the final submitted version.

## Extended data

**Extended Data Table 1.** Average recombination frequencies for the *attC* x *attI* suicide conjugation assays.

|  | *attC* bottom strand<br>(*intI1* induced*) | *attC* bottom strand<br>(*intI1* suppressed†) | *attC* top strand<br>(*intI1* induced) | *attC* top strand<br>(*intI1* suppressed) |
| --- | --- | --- | --- | --- |
| *attC<sub>aadA7</sub>* | 2.54x10<sup>-1</sup> | ND<sup>††</sup> | 2.48x10<sup>-3</sup> | ND |
| *attC<sub>Aenigmataarchaeota</sub>* | 5.46x10<sup>-4</sup> | ND | 8.66x10<sup>-7</sup> | ND |
| *attC<sub>Asgardarchaeota_1</sub>* | 1.79x10<sup>-2</sup> | ND | 6.74x10<sup>-4</sup> | ND |
| *attC<sub>Asgardarchaeota_2</sub>* | ND | ND | ND | ND |
| *attC<sub>Hadarchaeota</sub>* | 1.13x10<sup>-1</sup> | ND | 1.18x10<sup>-3</sup> | ND |
| *attC<sub>Halobacteriota</sub>* | 4.88x10<sup>-2</sup> | ND | 4.33x10<sup>-4</sup> | ND |
| *attC<sub>Hydrothermarchaeota</sub>* | 5.21x10<sup>-2</sup> | ND | 6.92x10<sup>-3</sup> | ND |
| *attC<sub>Nanoarchaeota</sub>* | 5.84x10<sup>-2</sup> | ND | 1.55x10<sup>-3</sup> | ND |
| *attC<sub>Thermoplasmatota</sub>* | 4.08x10<sup>-2</sup> | ND | 2.28x10<sup>-3</sup> | ND |
| *attC<sub>Thermoproteota</sub>* | 9.54x10<sup>-2</sup> | ND | 1.80x10<sup>-3</sup> | ND |

\*induced using L-arabinose; †suppressed using D-glucose; ††ND = Not detected

**Extended Data Fig.1:**
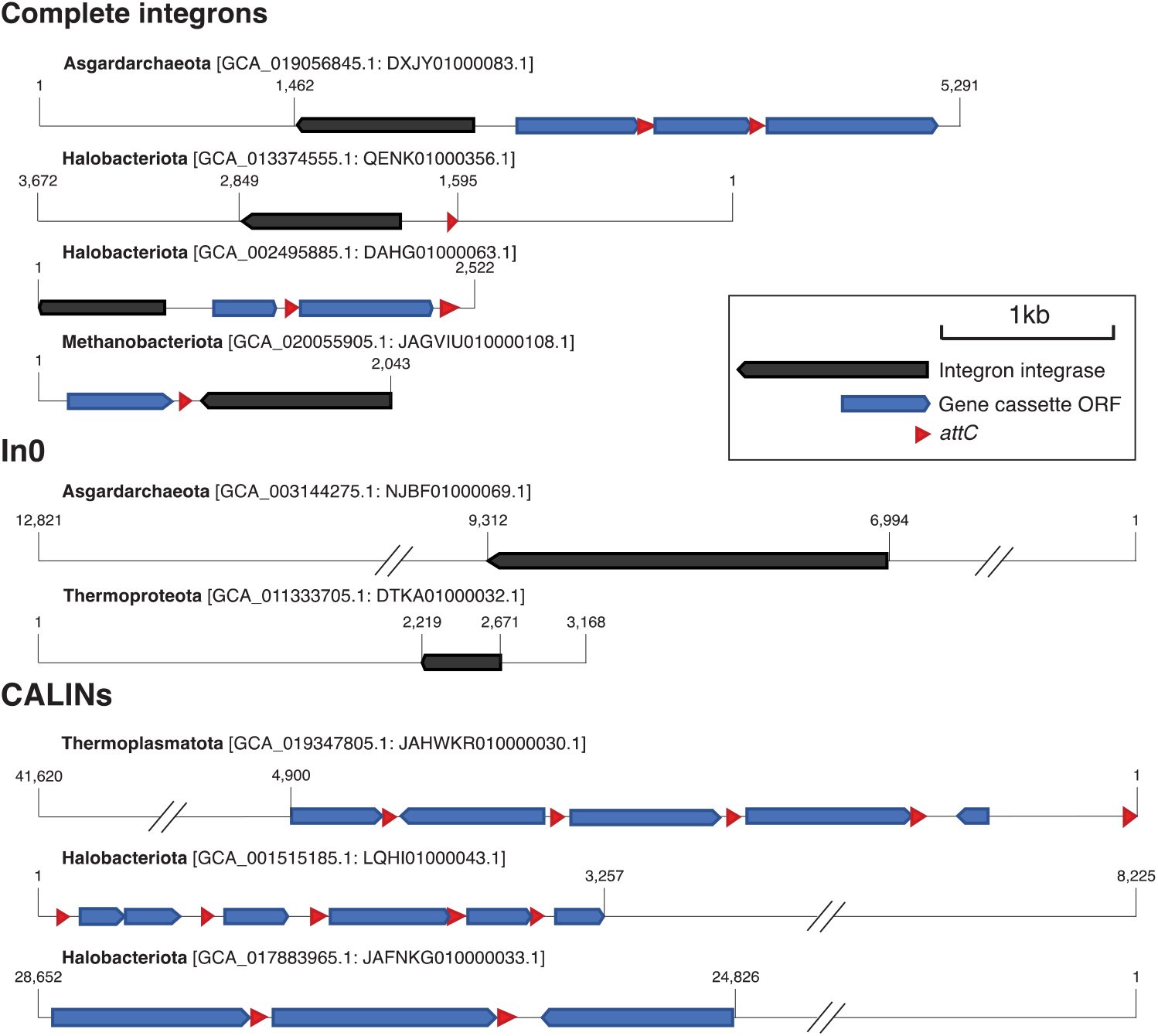
Example structure of archaeal integrons. Maps of all ‘complete integrons’, which are those that comprise an integron integrase gene (*intI*) and at least one gene cassette recombination site (*attC*); all ‘In0’ elements, which are those with *intI* but no detectable *attC* site; and three examples of ‘CALINs’ (clusters of *attC*s lacking integron integrases).

**Extended Data Fig.2:**
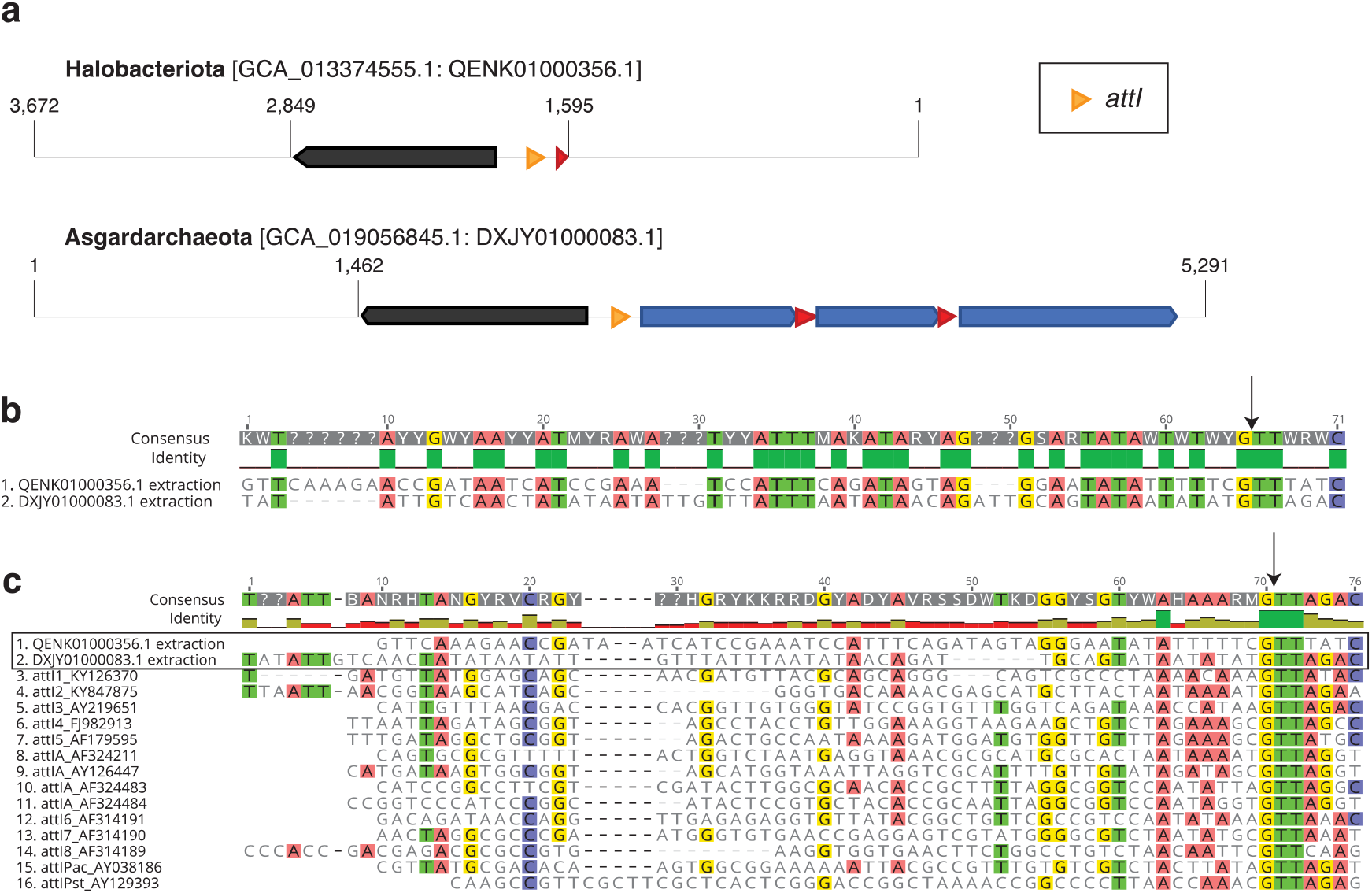
Putative archaeal integron recombination sites, *attI*s. **a**, maps showing the location of putative archaeal *attI*s. **b**, sequence alignment of the two putative archaeal *attI*s. **c**, multiple sequence alignment of the two archaeal *attI*s and all annotated bacterial *attI*s from the INTEGRALL database^80^. Nucleotides are coloured if they match with at least 50% of the sequences. Vertical arrows indicate the canonical insertion point of an inserting gene cassette.

**Extended Data Fig.3:**
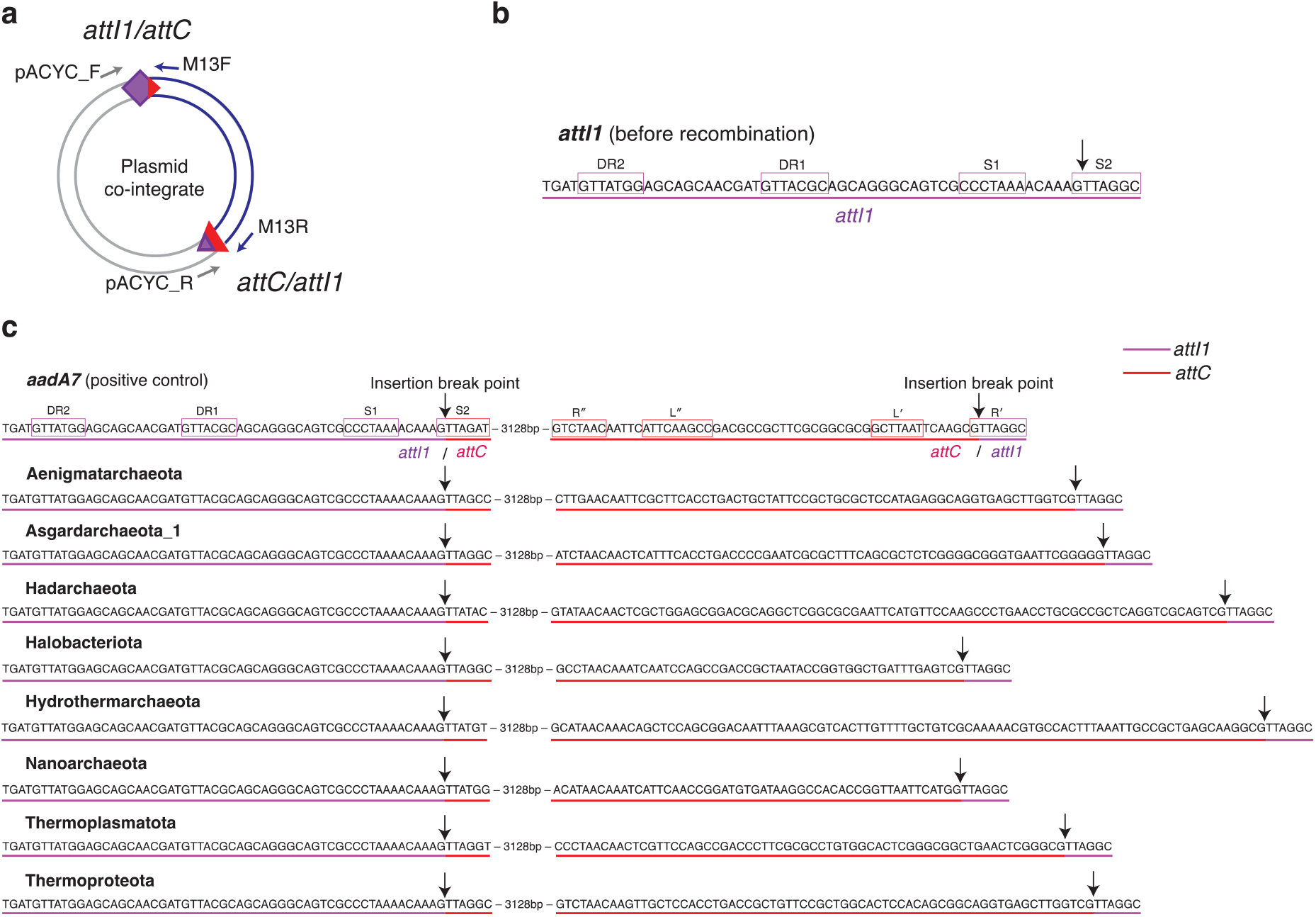
Sanger sequencing of *attI1* x *attC* recombination junctions. **a**, schematic of PCR primer pairs (grey and blue arrows) that amplify the recombination junctions following cassette insertion (*attI1* x *attC* recombination). **b**, *attI1* sequence before recombination. Boxes denoted with S1 and S2 indicate the core IntI1 binding sites, and the direct repeats signified by DR1 and DR2, are additional strong and weak IntI1 binding sites, respectively. The black arrow indicates the insertion break point where cleavage takes place during recombination. **c**, Sanger sequence data of the recombinant clones following *attI1* recombination with the paradigmatic bacterial *attC* site (*attC*_*aadA7*_), used as positive control, and eight archaeal *attC*s. Black arrows indicate the insertion break points following recombination. For *attC*_*aadA7*_, the two sets of paired inverted repeats are boxed (R′ to R″ and L′ to L″).

**Extended Data Fig.4:**
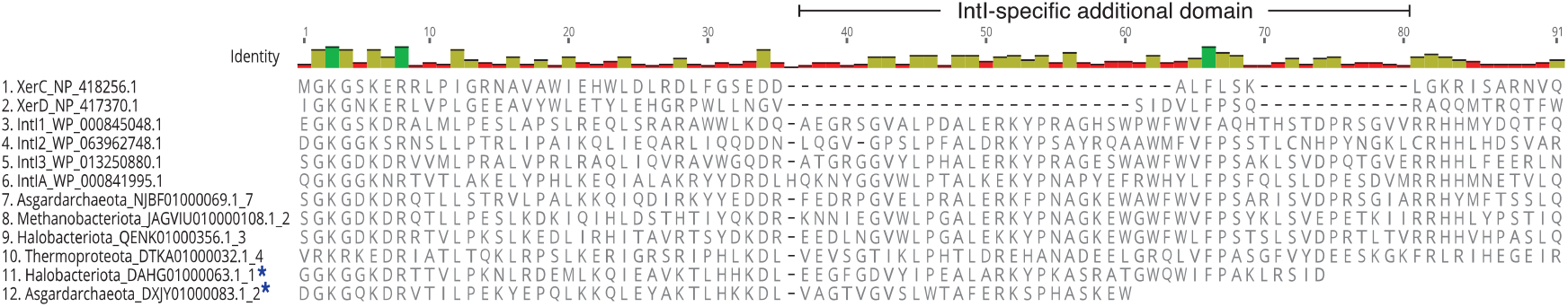
A multiple protein sequence alignment of the additional domain unique to integron integrases. Sequences (1) and (2) are tyrosine recombinases XerC and XerD that lack the IntI-specific domain. Sequences (3) to (6) are bacterial IntIs, and (7) to(12) are IntIs from Archaea. Blue asterisks indicate IntIs that did not span the full additional domain and were excluded from phylogenetic analysis.

**Extended Data Fig.5:**
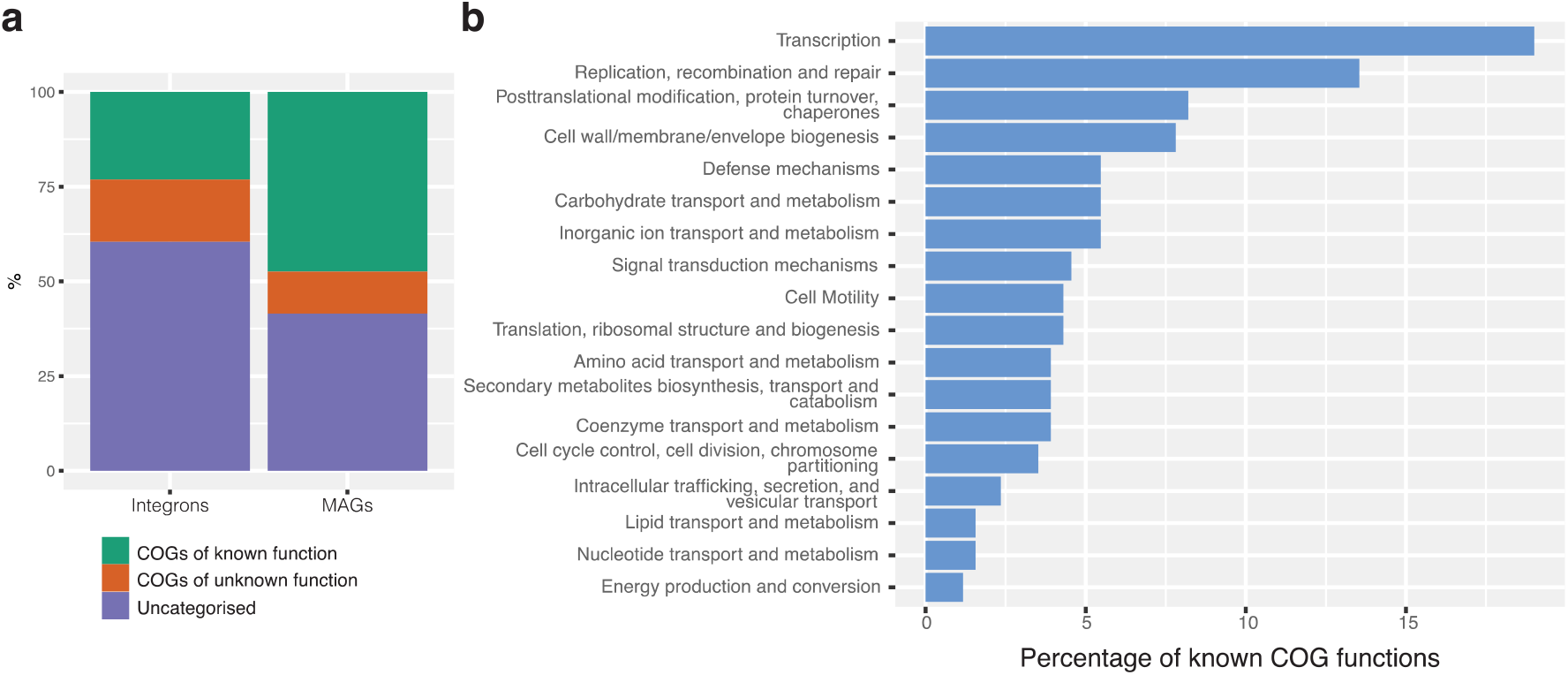
COG functional analysis of archaeal gene cassettes. **a**, percentage of proteins assigned a COG category. ‘Integrons’ represent all cassette-encode proteins in Archaea, while ‘MAGs’ indicate all proteins from the 75 integron-bearing archaeal genomes. **b**, percentage of COGs with known functions assigned archaeal cassette-encoded proteins.

